# The effects of repeated intraperitoneal administration of the GABA_B_ receptor agonist baclofen on body weight in rats maintained on a restricted feeding schedule

**DOI:** 10.1101/2023.01.24.525410

**Authors:** Ivor S. Ebenezer

## Abstract

It has previously been shown that chronic systemic administration of the GABA_B_ agonist, baclofen, reduces body weight in free feeding rats but has no effect on long-term food intake. The present study was conducted to extend these observations by investigating the effects of baclofen on body weight, food intake and water consumption in rats maintained on a restricted feeding schedule. The rats were allowed free access to food for 3 hours each day and injected intraperitoneally (i.p) once daily over a period of 11 days with baclofen (4 mg / kg; n=8) or saline (Control Group; n=8) immediately after each feeding session. Baclofen treatment produced small decreases in food intake on Days 2, 4 and 5 (P<0.05) compared with control animals but there were no differences between the 2 groups on Days 6 to 11. Baclofen did not affect prandial water intake. However, baclofen significantly decreased body weight gain of the animals (F_*(1,14)*_ = 22.0, *P* < 0.01) starting 3 days after the initiation of treatment. During the subsequent 9 days both groups received no treatments. However, the “Baclofen Group” continued displaying significant reductions in body weight gain (F_(1,14)_ = 8.47 p<0.01) but there were no significant differences in food and water consumption between the 2 groups of animals. The results of this study indicate that repeated administration of baclofen to food-restricted rats for 11 days reduces body weight and that the effects persist when the drug is withdrawn. While it is possible that the decrease in food intake during the early period of the experiment may have been partly responsible for the reduction in body weight in the baclofen-treated animals, the observation that these rats displayed reduction in body weight during the latter part of the study when there was no significant differences in food intake between the 2 groups, suggests that the drug decreases body weight independently of its effects on food consumption.

## 1. Introduction

It has previously been demonstrated in pig, rat and mouse, that central and systemic administration of the GABA_B_ receptor agonist, baclofen, increases food intake in satiated or non-deprived animals (Ebenezer and Baldwin, 1990; Ebenezer, 1990; Ebenezer and Pringle, 1992; Ebenezer, 1995; Ebenezer and Patel, 2004, Higgs and Barber, 2004; Buda-Levin et al., 2005; Ebenezer and Prabhaker, 2007; Patel and Ebenezer, 2010, 2008a and others) by a central GABA_B_ receptor mediated central mechanism of action (Ebenezer and Baldwin, 1990; Ebenezer, 1990; Ebenezer and Patel, 2004). More recently, it has been shown that repeated intraperitoneal (i.p.) administration of baclofen (1 and 4 mg / kg) once a day to non-deprived rats for 27 days also increased daily short term food intake, without tolerance occurring to the short term hyperphagia (Patel and Ebenezer, 2010). Repeated injections of baclofen had no effects on daily 24h food intake. Surprisingly, however, chronic treatment with the 4 mg / kg dose, but not the 1 mg / kg dose, of baclofen decreased body weight gain (Patel and Ebenezer, 2010). It was suggested that that the GABA_B_ receptor agonist may act through different mechanisms to influence food intake and body weight. One possibility that was mooted is that baclofen increases metabolic rate in a dose-dependent manner to decrease body weight gain (Patel and Ebenezer, 2010).

While most of the studies on the effects of baclofen on food intake have focussed on the effects of acute or repeated administration of baclofen on feeding responses and on body weight in non-deprived animals ((Ebenezer, 1990, 1995, 1996; Ebenezer and Pringle, 1992; Ebenezer and Patel, 2004; Patel and Ebenezer, 2008a,b, 2010; Ebenezer and Prabhaker, 2007; Bains and Ebenezer, 2013; Stratford and Kelly, 1997; Ward et al, 2000; Higgs and Barber, 2004; Buda-Levin et al., 2005), there has been a paucity of studies that have examined the effects of the drug in fasted animals.Ebenezer and Patel (2011) found that i.p administration of baclofen to 22h fasted rats had no effects on food consumption. However, when fasted animals were given a 2h oral preload prior to drug administration, baclofen (1 – 4 mg / kg) produced an increase in food intake. These authors suggested that the effects of baclofen on food consumption might be related to the state of hunger or satiety of the animals.

The present study was designed to extend previous observations obtained with baclofen by investigating the effects of repeated administration of the drug on body weight, food intake and water consumption in 21h fasted rats. In this study, baclofen was administered after each feeding session rather than prior to the feeding sessions, as reported previously in non-deprived animals (Patel and Ebenezer, 2008a, 2010). The effects on body weight, food intake and consumption were also recorded daily after drug treatment ceased to examine whether there were any long-term effects on these parameters.

## 2. Material and methods

The protocols used in this study were approved by the Ethical Review Committee at the University of Portsmouth, England and carried out under licences granted by the UK Home Office Scientific Procedures Act.

### 2.1 Effects of repeated administration of baclofen on body weight, food intake and water intake in fasted rats

Adult male Wister rats (n=16; starting body weights: 295 – 336 g) were housed in cages in groups of 4 where they had free access to water at all times. Male rats were used in this study so that the results could be compared with earlier studies in this area of research. The animals were maintained on a 12 h light / dark cycle (lights on at 8.30h and lights off at 20.30h) and were fasted for 22 h each day between 16.00 h –13.00 h. The rats were divided into 2 equal groups of similar body weights and were given 4 training sessions lasting 3 h when they were allowed free access to their normal food pellets (Food composition: protein 20%, oil 4.5%, carbohydrate 60%, fibre 5%, ash, 7% + traces of vitamins and metals) and water in experimental cages measuring 32 x 25 x 10 cm. The food was presented to the rats in shallow cylindrical cups, as described previously (Ebenezer, 1990). During the experimental sessions that followed, the rats were injected i.p. with either physiological saline solution (Group 1) or baclofen (4 mg / kg; Groups 2) at the end of each feeding session and replaced in their home cages. This was repeated on a daily basis over a period of 11 days. Both group of rats received no treatments after the feeding sessions in the experimental cages during the next 9 days.

Cumulative food and water intake were measured at the end of each feeding session as described previously (Ebenezer, 1990; Houston et al., 2011). Body weight for each rat was recorded each day between 12.00 h – 12.30 h.

### 2.2. Drugs

(±) Baclofen was purchased from Sigma Biochemicals, Dorset, UK. The drug was dissolved in physiological saline solution (0.9% ^w^/_v_, NaCl) to give an injection volume of 0.1 ml / 100 g body weight. Physiological saline solution was used in control experiments.

### 2.3. Statistics

The data from was analysed by 2 way analysis of variance (ANOVA) with repeated measures on time (days) followed by the Student-Newman Keul *post-hoc* test (Winer, 1971). As saline and baclofen were first administered after the feeding session at the start of the experimental period (designated as Day 0), for statistical purposes the effects of treatment on body weight and food and water intake were analysed for the data obtained on Days 1 to 11, and the effects of no treatment were analysed for the data obtained on days 12 to 20.

### 2.4. Body Weight Gain

The body weight data obtained for each rat were expressed as a percentage of the average animal’s body weight recorded on the day prior to and Day 0 of the experiment.

## 3. Results

### 3.1. Effects of repeated administration of baclofen on food intake, water intake and body weight in fasted rats

#### 3.1.1. Food Intake

The effects of baclofen (4 mg / kg, i.p.) or physiological saline on food intake recorded on Days 1 to 11 are illustrated in Fig. 1. Statistical analysis of the data showed that there were significant main effects of treatment (F_*(1,14)*_ = 7.11, *P* < 0.05) and time (days) (F_*(10,140)*_ = 10.7, *P* < 0.01), but no significant effects of treatment x time (days) interaction (*F*_(10,140)_ = 1.83, *NS*) *Post-hoc* tests showed that repeated treatment with baclofen produced small but significant decreases (at least P<0.05) in food intake on Days 2, 4 and 5 (Figure 2). However, there were no significant effects on food consumption between the 2 groups on Days 6 to 11.

**Figure 1.**
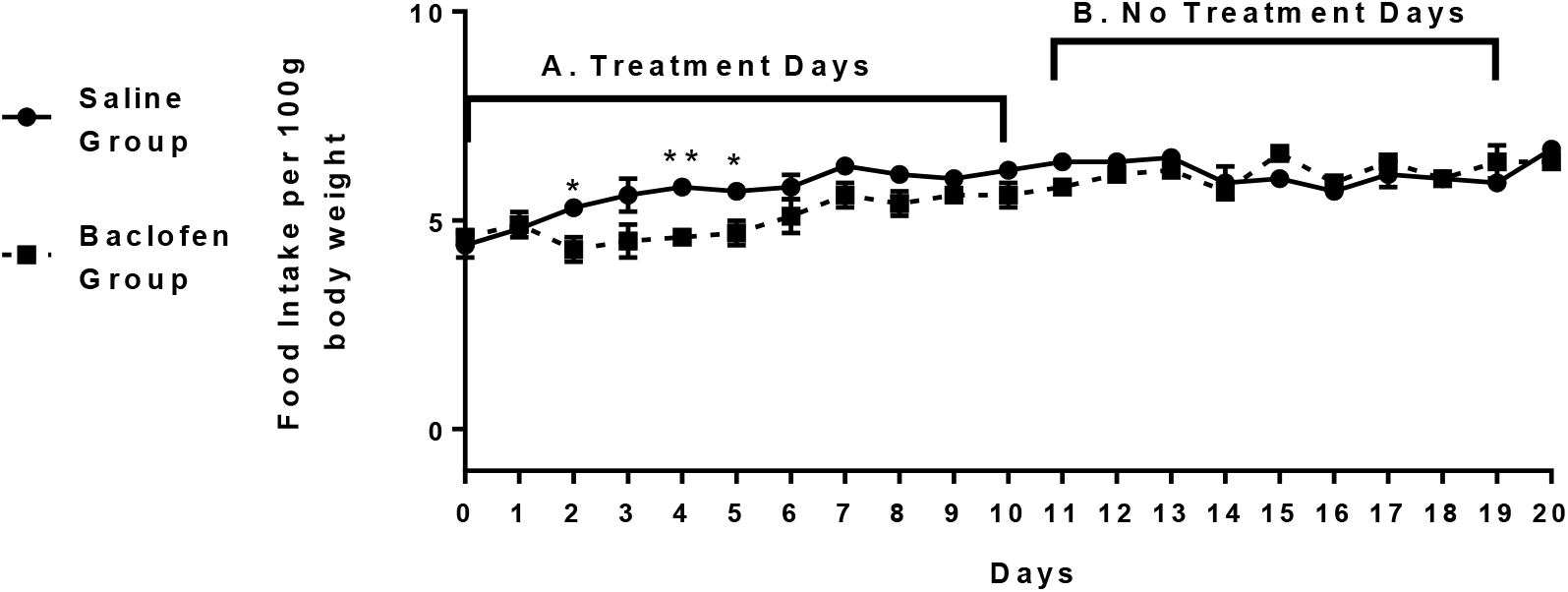
Effects of daily repeated intraperitoneal injections of physiological saline or baclofen (4 mg / kg) on 3h food intake in 21 h fasted rats. (N = 8 for saline and N = 8 for baclofen). A: Treatment Days; B: No Treatment Days. See text for further details. Vertical lines represent ±S.E.M. **P<0.01, *P<0.05

**Figure 2.**
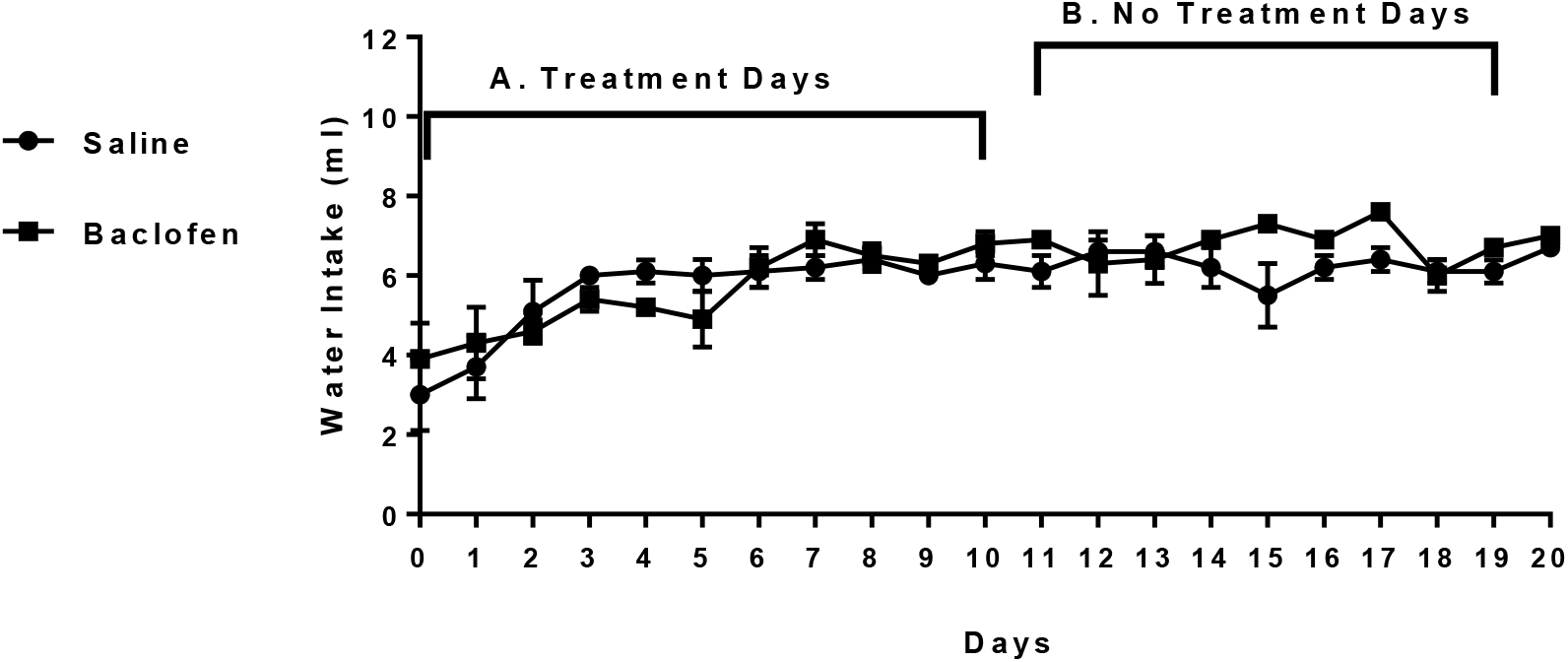
Effects of daily repeated intraperitoneal injections of physiological saline or baclofen (4 mg / kg) on 3h water intake in 21 h fasted rats. (N = 8 for saline and N = 8 for baclofen). A: Treatment Days; B: No Treatment Days.See text for further details. Vertical lines represent ±S.E.M.

After treatment ceased (Fig. 1), statistical analysis of the food intake data revealed that there were no significant effects on food intake between the two groups (*F*_(1,14)_ = 0.015, *NS*)

#### 3.1.2. Water Intake

The results are shown in Fig. 2. There were no significant effects on water intake between the two groups of rats either during the measurement periods when they were receiving repeated injections on saline or baclofen (F_*(1,14)*_ = 0.041, *NS*) or during the period when treatment had ceased ((F_*(1,14)*_ = 1.84, *NS*).

#### 3.1.3. Body Weight

The effects of baclofen (4 mg / kg, i.p.) or physiological saline on body weight gain after each feeding session during Days 1 to 11 are shown in Fig. 3. Statistical analysis of the results revealed that there were significant main effects of treatment (F_*(1,14)*_ = 22.0, *P* < 0.01) and time (days) (F_*(10,140)*_ = 10.9, *P* < 0.01), and significant effects of treatment x time (days) interaction (*F*_(10,140)_ = 7.58, *P* < 0.01) *Post-hoc* tests showed that there were significant reductions in body weight gain in the baclofen treated rats compared with controls animals starting on Day 3 and continuing until Day 11 (*P*<0.01 in each case).

**Figure 3.**
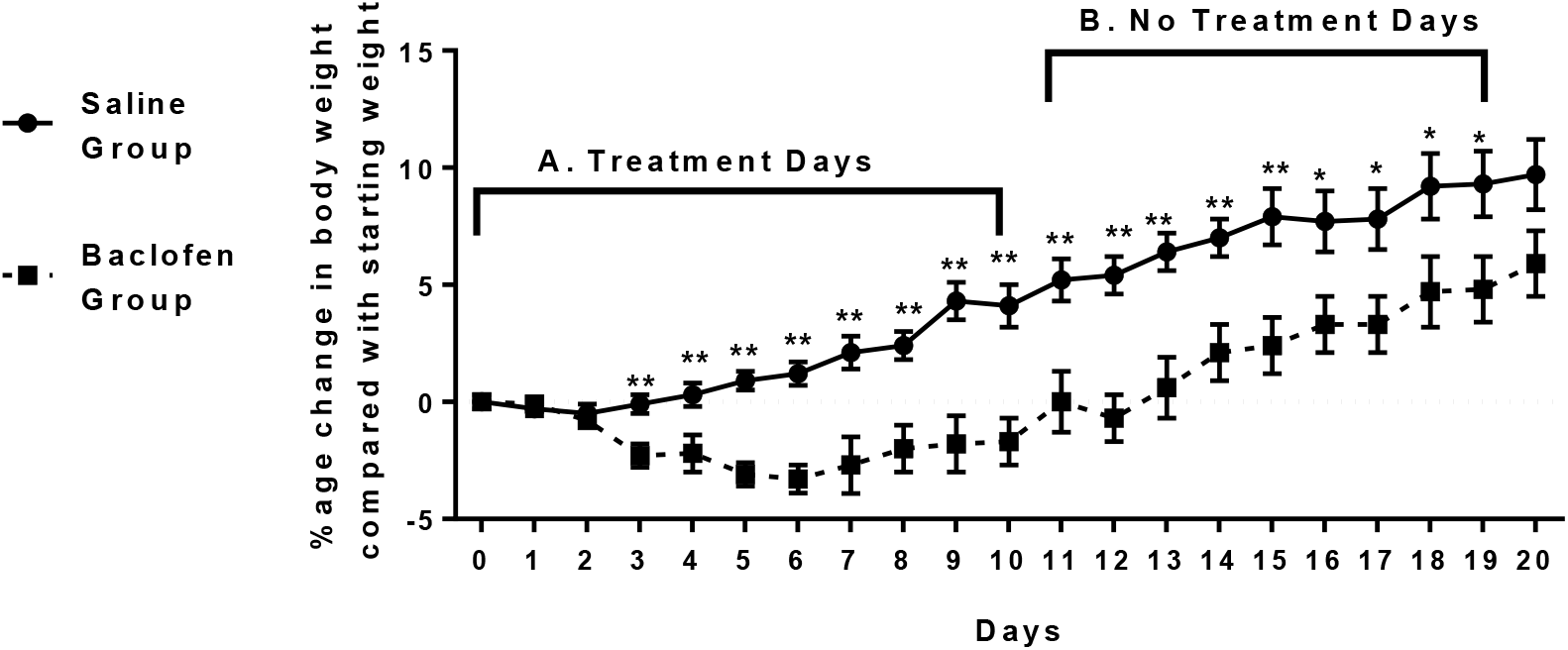
Effects of daily repeated intraperitoneal injections of physiological saline or baclofen (4 mg / kg) on body weight gain in 21 h fasted rats (N = 8 for saline and N= 8 for baclofen). A: Treatment Days, B: No Treatment Days. See text for further details. Vertical lines represent ± S.E.M. **P<0.01, *P<0.05

When treatment ceased (Fig. 3), statistical analysis of the body weight data indicated significant main effects of treatment (*F*_(1,14)_ = 8.47, *P* < 0.01) and time (days) (*F*_(8,112)_ = 46.1, *P* < 0.01), and significant effects on treatment x time (days) interaction (*F*_(8,112)_ = 2.11 *P* < 0.05). *Post-hoc* tests showed that there were significant reductions in body weight gain in the animals in Baclofen Group compared with those in the Saline Group from Days 12 to 19 (at least P<0.05). However, there was no significant difference between the two groups on Day 20.

## 4. Discussion

It has previously been reported that repeated administration of baclofen (4 mg / kg, i.p.) once daily for 27 days to non-deprived rats (a) increases daily short term food consumption, (b) has no effect on 24h food intake, and (c) reduces body weight gain (Patel and Ebenezer, 2010). The present study was undertaken to extend previous observations and investigate the effects of repeated administration of baclofen (4 mg / kg) on body weight, food intake and water consumption in 21 h fasted rats when administered after each daily feeding session.

Baclofen (4 mg / kg) administered after each 3 h feeding session in 21 h fasted rats during the 11 day post-injection trials produced small, but significant, decreases in food intake on Days 2, 4 and 5 but there was no significant differences between the 2 groups on Days 6 to 11 (Fig. 1). It was demonstrated previously that administration of baclofen (4 mg / kg, i.p.) had no effects on food consumption compared with saline injection in 22 h fasted rats when the animals were given access to food immediately after saline or drug administration (Patel and Ebenezer, 2011). It was suggested that, while baclofen increases food intake in non-deprived and partially satiated animals (Ebenezer and Pringle, 1992; Ebenezer and Patel, 2011), hungry animals will eat maximally and therefore the effects of a drug that stimulates feeding in non-deprived animals may not be apparent in fasted animals (Ebenezer, 1990, Ebenezer and Patel, 2011). It is not clear why there was a small but significant decrease in food intake in some of early feeding trials in the present study. We have previously demonstrated that this dose of baclofen was not aversive to rats in a 2-bottle taste aversion test (Ebenezer et al, 1992). However, it is noteworthy that in this study baclofen was administered immediately after each feeding session and it is possible that the drug may have had some adverse effects, which, in turn, may have affected food consumption in early trials. It is well documented that this dose of baclofen can cause ataxia, muscle relaxation and sedation (Ebenezer and Pringle, 1992; Patel and Ebenezer, 2008a,b, Patel and Ebenezer, 2010, Bains and Ebenezer, 2013). The animals may have associated eating the food in the experimental cages with these effects of the drug and therefore ate less food during the early feeding trials (see Wilson et al., 2011). However, tolerance quickly develops to these effects of the drug (Patel and Ebenezer, 2010; Bains and Ebenezer, 2013) and this may explain why in the later feeding trials there were no effects on food intake between the two groups of rats. It is also unlikely that baclofen would have had a direct effect on feeding mechanisms as it has a half-life of approximately 4 h (Popova et al., 1995) and food was presented to the animals 21 h after administration..

When the treatments were discontinued (see Fig.1), there were no significant differences in daily food consumption between the two groups of rats. The amount of food consumed by the animals in each group when treatment ceased was very similar to that recorded during Days 6 to 11 when the animals were injected daily with saline or baclofen.

It has previously been shown that systemic administration of baclofen inhibits both osmotic and volemic drinking in rats by GABA_B_-receptor mediated mechanisms of action (Ebenezer et al., 1992; Huston et al., 2012). It has also been found that baclofen inhibits prandial drinking (Ebenezer and Baldwin, 1990; Ebenezer and Pringle, 1992). In the present study, there were no effects on the baclofen treated rats compared with the control rats on prandial drinking (see Fig. 2). These results were not unexpected, given that baclofen has a half-life of 4 h (Popova et al., 1995) and water intake was measured over a 3 h period 21 h after administration of the drug.

The rats used in this study were young adult rats and under normal circumstances they would continue to grow and display increases in body weight. The data obtained show that baclofen (4 mg / kg) reduced body weight gain compared with control animals (see Fig. 3). Significant reductions in body weight gain were apparent after the 3rd injection of baclofen and persisted throughout the treatment period. It is noteworthy that for most of the treatment period the body weights of the baclofen treated rats were below their pre-treatment starting weights (see Fig. 3). It is possible that the small decreases in food intake noted during the early period of the experiment (Days 2, 4 and 5; Fig.1) may have been partly responsible for the reduction in body weight in the baclofen treated animals. However, the observation that the baclofen treated rats still displayed significant weight loss compared with saline treated animals during the latter part of the study (Days 6 to 11; Fig. 1) when there was no significant differences in food consumption between the 2 groups, suggests that the early reductions in food consumption were probably not responsible for the weight loss

The body weight data are consistent with results obtained previously in which it was found that chronic administration of a 4 mg / kg dose of baclofen to free feeding rats significantly reduced body weight gain (Patel and Ebenezer, 2010). In that study, Ebenezer and Patel (2010) demonstrated that while daily dosing with baclofen reduced body weight gain, it did not affect 24 h food intake. The results indicated that the inhibitory effect of the GABA_B_ receptor agonist on body weight gain is not related to long-term suppression of food intake. The authors suggested that it was likely that the effects of baclofen on food intake and body weight may be mediated by separate mechanisms. It was proposed that baclofen might reduce body weight by increasing metabolic rate by acting in the ventromedial nucleus of the hypothalamus (VMH) to increase activity of the sympathetic nervous system to activate brown fat (see Patel and Ebenezer, 2010; Rothwell et al; 1985; Addae et al., 1986). However, it is also possible that other mechanisms may be involved to increase metabolic rate. Interestingly, it was found that even when daily injections of baclofen were stopped, the rats still displayed significant reductions in body weight gain compared with the control animals (see Fig. 3). These data suggest that the GABA_B_ receptor agonist may initiates changes in brain biochemistry that result in a protracted suppression of body weight during drug treatment and long after withdrawal of the drug by increasing metabolic rate, or by some other mechanism(s).

In conclusion, the main findings of the study extend previous observations (Patel and Ebenezer, 2008a, 2010, Ebenezer and Patel, 2011) and show that repeated intraperitoneal injections of baclofen administered after each feeding trial in 21 h fasted rats (i) elicits a small hypophagic effect during some early but not later feeding trials when the animals are given access to food for 3 h, (ii) has no effects on prandial water intake, and (iii) reduces weight gain that persist even when the drug is withdrawn. While it is possible that the decrease in food intake during the early period of the experiment may have been partly responsible for the reduction in body weight in the baclofen-treated animals, the observation that these rats displayed reduction in body weight during the latter part of the study, when there was no significant differences in food intake between the 2 groups, indicates that the drug decreases body weight independently of its effects on food consumption. Furthermore, the effects persisted for many days after the drug was withdrawn and implies that baclofen initiates long-term changes in brain systems that result in continued suppression of body weight. These data suggest the possibility that GABA_B_ agonists, such as baclofen, may be useful in the treatment of obesity. In support of this suggestion, a preliminary open label, single arm, clinical trial has indicated that baclofen reduces body weight in obese human subjects (Arima and Oiso, 2010). However, further work needs to be undertaken to investigate the mechanisms involved in mediating the effects of baclofen on body weight and to validate the preliminary clinical data.

